## Supplementary material for "Comparing a computational model of visual problem solving with human vision on a difficult vision task": S1 Text

**S1 Text. Adaptation of GenSearch on more complex THINGS dataset** We also tested the model on the more complex constellations created using the Things dataset [1]. As the model needs to work with a more general set of things, i.e. 1100 categories, we use stable diffusion [2] as the generator to sample the images. In the stable diffusion model, search is done on the encoded latent space of the language model that conditions the image generator. In this case, the search algorithm fails to find the relevant solution category using the search and fit strategy starting from a random population. Until the end of max iterations, the search mostly cannot converge to a single solution. The analysis of the solving process shows that the detailed nature of features generated by the stable diffusion generator causes spurious curves to match some dot features in the image and distract the search process (see. Fig. 1.a). We introduced a bottom-up initialisation for the initial population by finding straight lines and smooth curves (dots connected by lines with angles less than 10 degrees). This allowed the model to find the solution when such lines and curves made prominent structures of the object (example in Fig 1.b). The overall accuracy of this model on the dataset is 18% compared to the human performance of 65%. This modification and the nature of failure indicate that better initialisation and a model to generate the object outlines at the correct level of abstraction are required to improve the models to work on more general constellation object datasets.

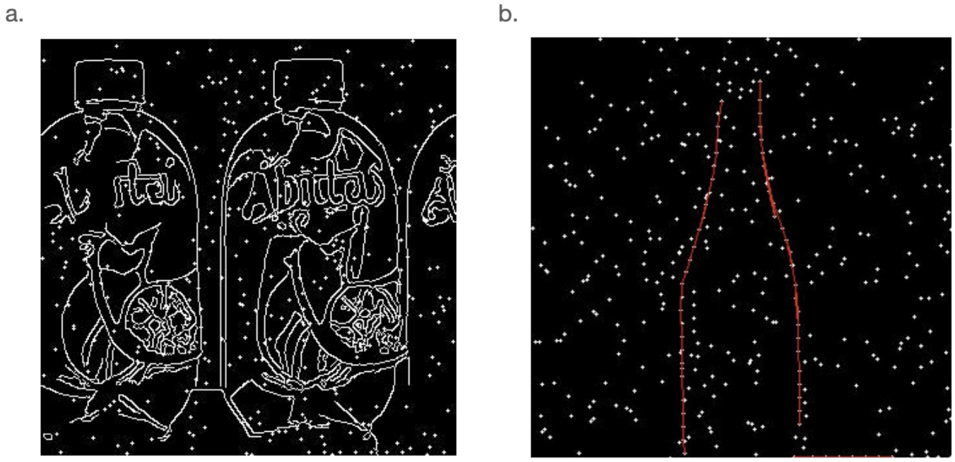


**Fig 1. GenSearch on THINGS constellations** a) GenSearch fails to work well, in its original form, on the more general THINGS constellations dataset mostly due to the complex and spurious patterns, such as design and packaging labels generated by the stable diffusion generator, which do not define the object b) adding a bottom-up prior, to identify straight lines and curves before initialising the search helps improve performance.
