## Supplementary material for "Comparing a computational model of visual problem solving with human vision on a difficult vision task": S2 Text

**S2 Text. GenSearch's convergence vs human reaction time** Another way we compared the process followed by GenSearch to the human process of solving the images is by evaluating the time taken to solve across an image set. The time taken for GenSearch is approximated by the convergence iteration (or generation). We do not see a clear correlation between the datasets and average human reaction time in this comparison 1. However, we also note a quite high variance in human reaction time to the images, making it difficult to use the relation to measure the model's quality confidently.

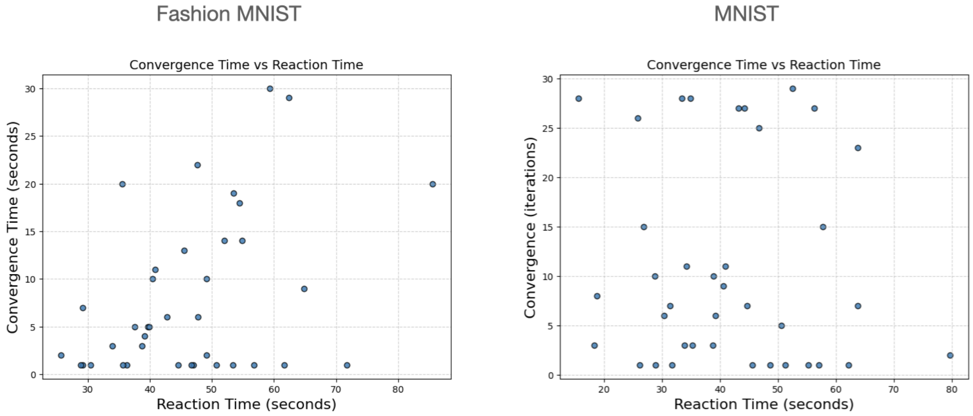


**Fig 1. GenSearch's convergence vs human reaction time** We plot the number of iterations used by GenSearch vs average human reaction time on the images for the datasets. There is a slight correlation (Pearson's coefficient =0.4096, pvalue=0.010). between these two measures for Fashion MNIST, but for the MNIST, they are uncorrelated (Pearson's coefficient =-0.072, pvalue=0.66). We note that the noise ceiling for this correlation is not high, i.e., we see a low left-out correlation agreement between participants and other participants (0.21 for Fashion MNIST and 0.35 for MNIST).
