## Supplementary material for "Comparing a computational model of visual problem solving with human vision on a difficult vision task": S3 Text

**S3 Text. Additional analysis of Resnet18 models** We use Grad-Cam, a method to analyse the gradient projection with the resulting class on the feature maps at various layers. These are then aggregated into a heatmap representing the contribution of areas of the image to the decision taken by the classifier. As we can see in Fig. 1 the highlighted area in the image does not project to the overall shape of the object or any distinguishing feature of the object class. Rather, over the images, we see that the classifier learns to pay attention to the presence (density) of dots in a part of the image for a particular category.

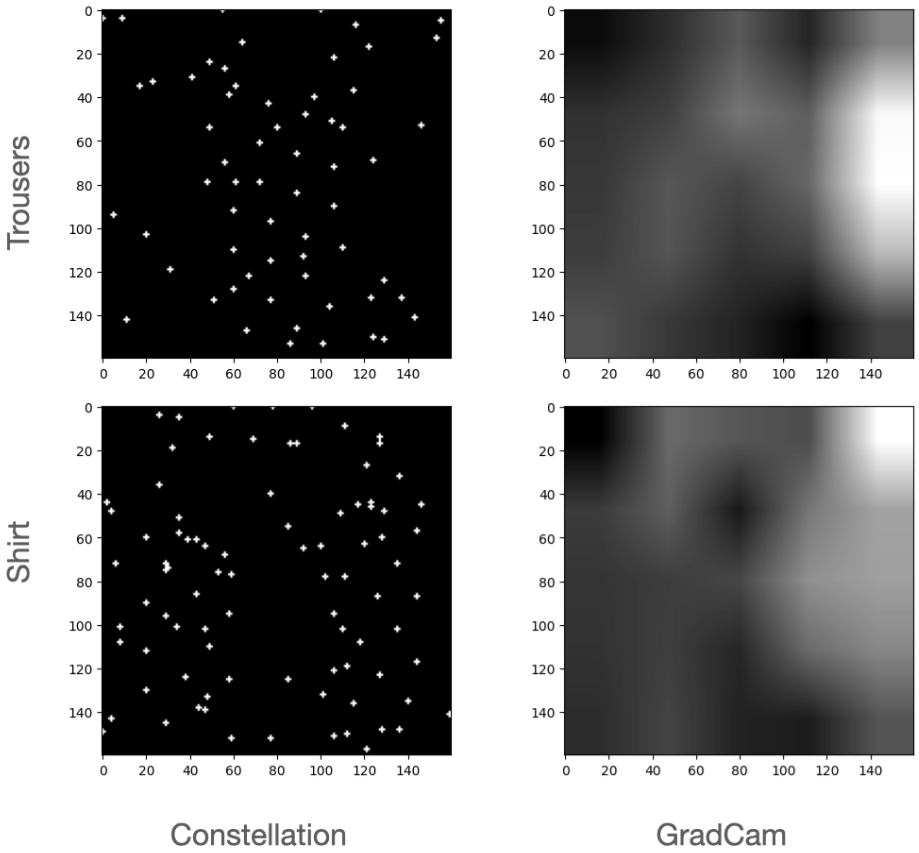


**Fig 1. GradCam result for Resnet18** The figure shows the constellation images along with their Grad-Cam projection for the prediction by Resnet18 on the image. Both images have been classified correctly by the trained Resnet18 model as Trousers and Shirt, respectively. The Grad-Cam projections, however, point out that the areas contributing the most are empty spaces in the image, hence showing Resnet's greater reliance on learning and using the distribution of dots in the image space for identifying the class rather than using the shape or feature of the object.

Further, we evaluated the confusion matrix of the models trained on MNIST and Fashion MNIST constellations datasets on the test set and report the confusion matrix in 2. The correlation of these matrices to the human confusion matrix is 0.68 for

Fashion MNIST and 0.66 for MNIST, which is lower than the correlation of GenSearch's confusion matrices. As these models have very high accuracy, the correlation captures the corresponding correct solution categories, but the mistakes made by humans are not captured by these models.

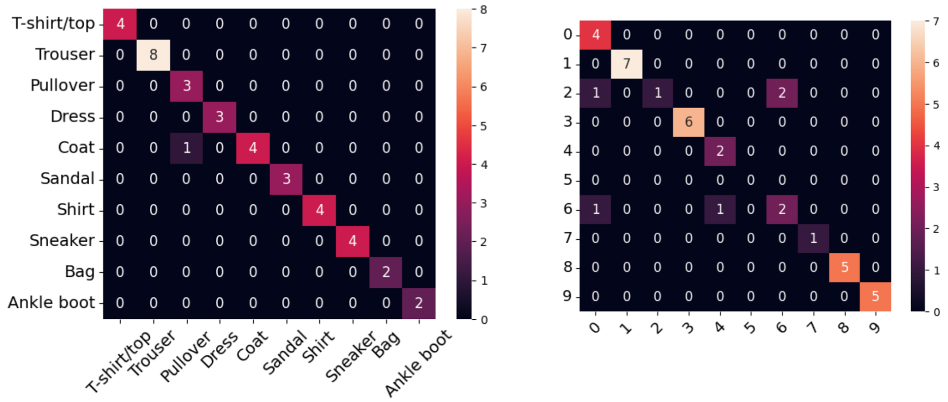


**Fig 2. Confusion Matrix for Resnet18 on 38 test set images** The figure shows the confusion matrix for Fashion MNIST (left) and MNIST (right) datasets evaluated by classifying these sets using their respective Resnet18 models trained on constellation images.
