## Supplementary material for "Comparing a computational model of visual problem solving with human vision on a difficult vision task": S4 Text

**S4 Text. Pix2pix training setting and hyperparameters search** All pix2pix models were trained on the pair of original and constellation train sets of the respective datasets. In Table 1, we show the test set performance for the variation in initial learning rate while using the linear decay schedule. We also tried other learning rate schedules such as cyclic and reduce on plateau, but the linear decay worked the best and is also reported to be working best with models such as Pix2Pix.

**Table 1. IOU dots with correct solution for tuning initial learning rate for pix2pix on both datasets**

| Learning Rate | MNIST | Fashion MNIST |
|---|---|---|
| 0.1 | 0.0 | 0.0 |
| 0.01 | 0.38 | 0.18 |
| 0.001 | 0.30 | 0.23 |
| 0.0001 | 0.30 | 0.23 |
