## Supplementary material for "Comparing a computational model of visual problem solving with human vision on a difficult vision task": S5 Image

**S5 Image. Examples of images generated by GANs**

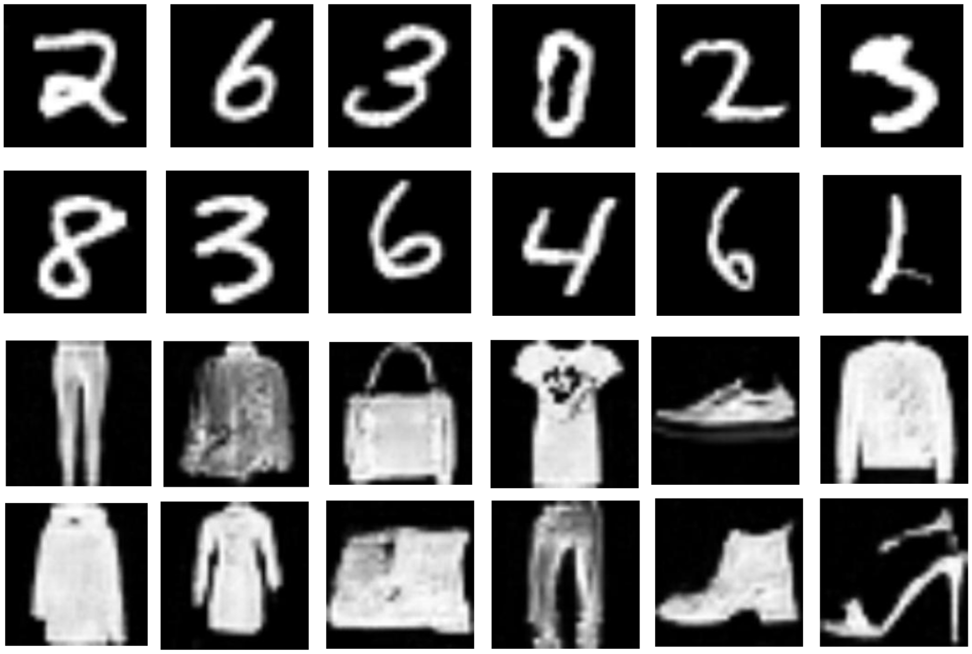

**Fig 1. Examples of images generated by the GANs** The figure shows images sampled from the GANs used in the GenSearch algorithm.
